# PROTEORIZER: A holistic approach to untangle functional consequences of variants of unknown significance

**DOI:** 10.1101/2024.07.16.603688

**Authors:** Torsten Schmenger, Gaurav Diwan, Robert B. Russell

## Abstract

Most *in silico* tools only use data closely related to the gene-of-interest or initial research question. This gene-focused research is prone to ignoring low-count and rare variants in the same or similar genes, even if available informational could be sufficient to deduce functional consequences by combining knowledge from many similar genes.

Proteorizer is a web tool that aims to bridge the gap between protein-centric knowledge and the functional context this knowledge creates. We use curated and reviewed data from UniProt to collect available residue information for the queried protein as well as orthologs. By defining functional clusters based on intramolecular distances of residues with available functional information it is possible to use these to extrapolate the effect of a VUS solely based on known functions of nearby residues, hence contextualizing the variant with pre-existing knowledge. We show that pathogenic variants are more likely to be a part of functional hotspots and present several case studies (ALPP p.Ser244Gly, CANT1 p.Ile171Phe, ARL3 p.Tyr90Cys, IL6R p.His280Pro and RAF1 p.Ser259Ala) to highlight the applicability and usefulness of this approach.

**Graphical Abstract:** 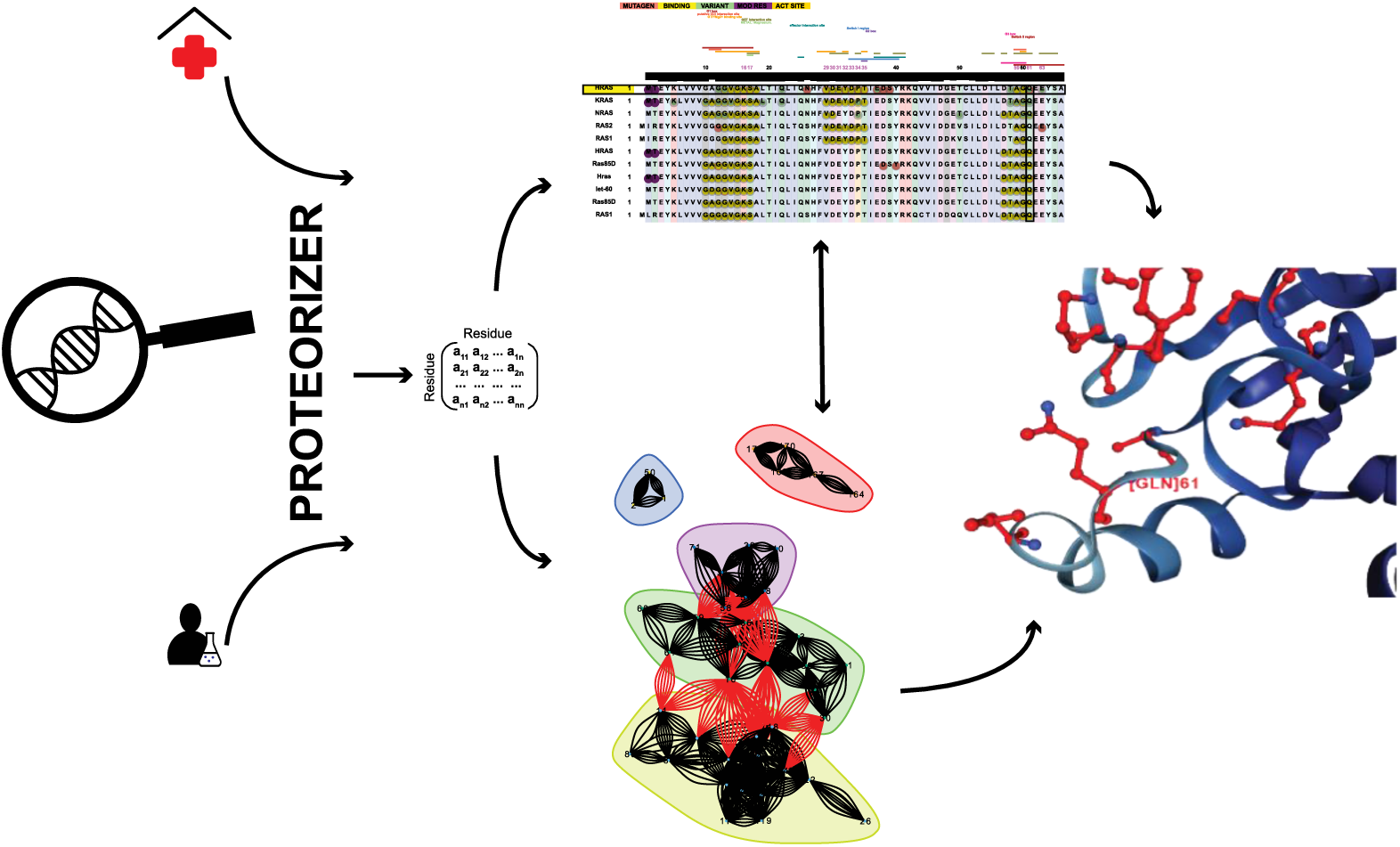

**Proteorizer** is an explorative tool that takes variants from laboratory or clinical settings and contextualizes the variants based on prior information from the protein of interest and similar proteins according to where these functional positions are located in the 3D structure of the protein of interest.

## Introduction

Despite tremendous advances in sequencing technologies and data generation the understanding of novel missense variants remains a difficult and challenging task. Millions of missense variants in gnomAD^1^ currently lack a functional assessment, despite the possibility that many might affect protein function^2^. Meanwhile, the vast majority of human proteins are sequenced^3^, and while experimental 3D structures are not yet available for every protein, many thousands of 3D structures^4^ are available to the public and structural gaps can be filled with reasonable predictions thanks to AlphaFold^5^.

Previous research in this field highlighted how protein structures can be used to predict the impact of missense variants of protein interactions^6^, and structure analysis proved useful in identification of mutational hotspots^7^, suggesting that such regions become hotspots for mutations due to their importance to protein function. Since then, several approaches have been made to use 3D structure constraints to predict the impact of variants of unknown significance (VUS)^8–10^, with promising results^11^.

Most *in silico* tools however only use specific data closely related to the gene-of-interest and to the initial research question. This gene-focused research often is conducted in an artificially narrow scope by ignoring low-count or rare variants in the same or similar genes^12^, even if the informational coverage (any functional information of the gene of interest) could be significantly increased by combining a variety of information from different but similar genes and datasets^13^.

Here, we present Proteorizer (shiny.russelllab.org/proteorizer), a new hypothesis and web tool that aims to bridge the gap between the available and increasing protein-centric knowledge and the functional context this knowledge should in principle create (but often doesn’t). Proteorizer uses the curated and reviewed data from UniProt^14^ to collect available residue information for the queried protein as well as orthologous proteins. Proteorizer also forms functional clusters based on the intramolecular distances of residues with available information using Alphafold2 structures. These clusters can then be used to extrapolate the effect of a VUS based on known functions of nearby residues, hence contextualizing the variant with pre-existing knowledge. We offer different selections of information data sources and methods for intramolecular clustering.

## Material and Methods

### Web Implementation

The online version of Proteorizer is based on the Shiny framework^15^ for development of web-based applications using the R programming language^16^. Tables are visualized with DT^17^ and AlphaFold structures are displayed using NGLVieweR^18^.

User queries are processed by both R and Python 2.7 and then sent to a Postgres database^19^. Afterwards, the proximity clustering is performed in R (see below) and results are generated. The annotated alignment^20^ is created in python using svgwrite^21^. All scripts are available in the GitHub repository (https://github.com/tschmenger/PROTEORIZER) and a general alignment annotation script is offered as a standalone version on python 3.6. (https://github.com/tschmenger/Annotate_Alignments).

The web app is divided into multiple sections including the landing page ‘Introduction’, containing instructions on how to format the input and some background on the method and formatting of custom alignments. The ‘Submit’ section offers different cluster methods and functional information datasets. Variant searches do not require a signup or login and while the user can provide custom alignments users must only provide a gene name or uniprot accession and a variant (“P04637/R273C” or “EGFR/L858R”) to use the tool, and example submissions are provided. The ‘Explore the results’ page displays the generated data. All data successfully queried by the user can be downloaded (data table, cluster plots, annotated alignment(s) and structure snapshots). The ‘Discover’ section provides access to two browsable datasets with contextualized VUS taken from humsavar (last updated September 2023).

### Clustering based on intramolecular distances and the detection of functional hotspots

Available positional information for a protein is selected from UniProt^14^. Sets of orthologous proteins are selected from pre-computed alignments generated through OrthoFinder^22^. These alignments are used to define the degree of sequence conservation for each position in the protein of interest and to map back functional information back to the query.

We defined functional residues as positions associated with a disease (‘VARIANT’) or known protein function (‘MOD_RES’, ‘ACT_SITE’ or ‘BINDING’). Additionally, we also considered experimental results (‘MUTAGEN’).

Collected Alphafold^5^ structures were used for distance measurement. Intramolecular (using C_α_ for glycine and C_β_ for all other amino acids) atomic distances between all *functional*-*residues* are measured and the average is kept. To avoid measuring distances between sidechains on opposing sides of the backbone (which would point away from each other) we compared two distances: residue-1-Cα vs. residue-2-Cβ to the default measurement of residue-1-Cβ vs. residue-2-Cβ (ignoring this comparison for Gly) and kept the shorter distance value (**Fig. 1C**). This captures extreme cases where sidechains of a given pair of amino acids are very unlikely to interact. Atomic distances ≤ 8 Å (**Fig. 1B**) are then subjected to either a random walk algorithm^23^ or hierarchical clustering^24^ to define clusters of putatively functional residues, based on intramolecular distances (**Fig. 1A**).

**Figure 1.**
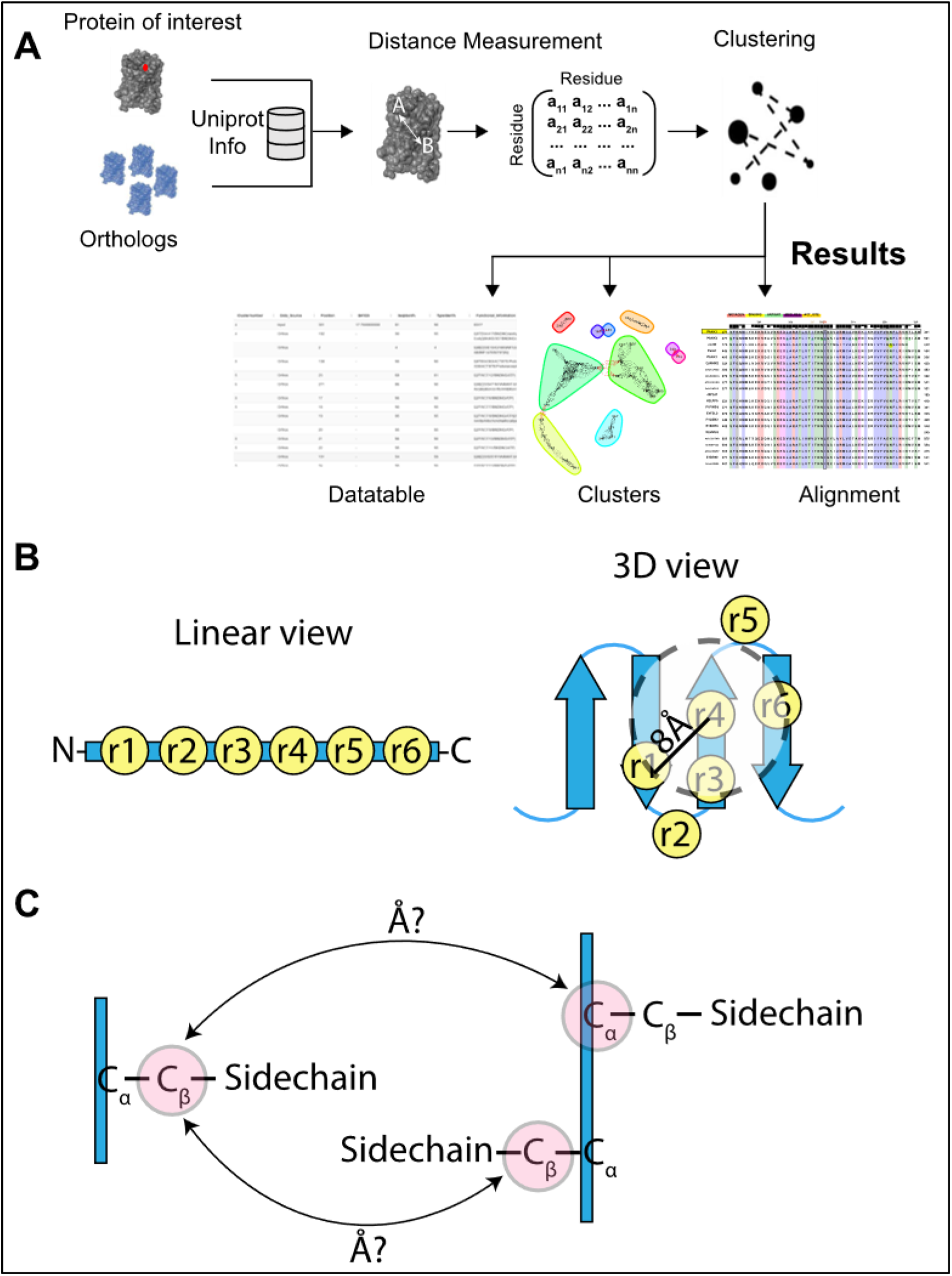
Workflow overview. **A.** General overview. Information on the protein of interest and orthologs is taken from Uniprot and mapped back to the queried protein. The distances between residues with functional information is used for clustering and results are generated. **B.** The Linear view (left) shows 6 residues (r1 to r6) with functional information. Only the 3D view (right) is able to group residues together based on where they are positioned on the protein structure. **C.** Distances are measured from Cβ to Cβ where possible and without violating the backbone.

### Creating a Bayesian evidence score

In addition to considering sequence conservation (identity & physicochemical properties) we created a feature vector containing counts for the total number of clusters and the total count of information we collected or mapped back to the queried protein. Additionally, if the position of interest was part of a cluster, we considered the number of residues in the respective cluster and the information content of that cluster as features, while removing available information on the queried position itself. We then combined these counts into a score using Bayesian integration^25,26^.

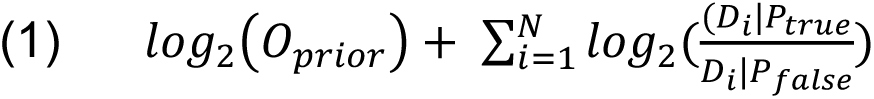

Where *D_i_*|*P_true_* and *D_i_*|*P_false_* correspond to the true and false positive rates (TPR and FPR), which were obtained from ROC curves considering 32385 known disease causing variants from Humsavar^14^ as positives and 39611 as negatives. We set *O_prior_* = 1 arbitrarily^2^.

### Developing a combined predictor

To support the naïve Bayesian classifier to distinguish likely functional variants from neutral variants we constructed an ensemble of decision trees using a random forest algorithm (Scikit-learn^27^, python 3.6) for both clustering methods (RW and HClust, **Supplemental Figure 4A-D**).

10-fold cross validation was performed to tune the parameters of the predictors including the number of decision trees in the forest (best: 78 for RW and 96 for HClust) and maximum tree depth (best: 10 for RW and 13 for HClust) and repeated the procedure 40 times. We evaluated the Accuracy (0.85 for RW, 0.91 for HClust), Precision (0.88 and 0.91), Recall (0.83 and 0.89) and Receiver Operator Curve (ROC, 0.92 and 0.96, (**Supplemental Figure 4C**) for both predictors and used the feature_importances_ function (Scikit-learn^27^) to determine the respective importance of all used features (**Supplemental Figure 4B**). We combined the predictions made by these predictors with the respective naïve Bayesian classifier and tested our approach on a small set of 11 variants with known functional consequences (neutral or not neutral) that were not part of the training set (**Supplemental Figure 4D**).

Due to the reliance on prior knowledge a ‘No Impact’ prediction based on absence of any known functional information has to be carefully evaluated by the researcher and we recommend using this prediction score as additional functional evidence but not as a final assessment (see Results and Discussion). To better reflect this limitation, we conservatively transformed the scores derived from both methods: Values from 10 to 15 were transformed into values ranging from 0.125 to 0.375 for the naïve Bayesian classifier and we likewise transformed scores between 0.5 and 1 for the random forest predictor. Only combined scores of 0.25 or higher were considered as predicted to have a functional impact (0.25 – 0.5 as ‘medium’ and > 0.5 as ‘high’). Both the combined verdict and the individual scores are displayed in the web app.

## Results and Discussion

### Identification of functional hotspots

Sequence analysis has been widely used for decades to study protein function and elucidate putative importance of particular residues, for example based on their conservation among similar proteins across different species. This approach is now revitalized and revolutionised by large-scale machine-learning^28^. Recently developed tools approached the challenge to explain and predict consequences of VUS based on prior-knowledge, for example assessing putative PPI consequences (Mechismo^6^) by using homologous structures or by summarizing available phenotypic and functional knowledge (Mechnetor^29^). While these and other tools generally share their use of available information to formulate new hypotheses, they rarely make use of information outside what would be seen as their research interest (in particular proteins).

The here presented approach makes use of sequence analysis to define a set of orthologous proteins^2^. This set of sequences is not only used to determine sequence conservation, it is also used to retrieve functional information which is mapped back to the protein of interest. A similar approach focusing on protein kinases has recently been shown to yield promising results^20^. For this study, we considered several different types of UniProt information (binding, active site residues, observed disease variants, post-translational modifications and mutagens). Grouping residues where we have direct functional knowledge for the protein of interest with known functions from orthologs based on their intramolecular distances can help elucidate the impact of VUS, offering guidance as to why particular phenotypes may be observed.

Applying this approach to a test set of benign and disease-causing mutations shows that proteins with disease-causing variants are more likely to produce functional hotspots, and such variants of interest are more often part of a functional cluster if the mutation is disease-causing (p < 1×10^−16^, **Supplement Figure 1**) regardless of the clustering method used. A scoring approach using a naïve Bayesian classifier combined with a random forest predictor further indicates that a variant being part of such a functional cluster may have putative functional consequences (**Supplement Figure 2**). We also tested this workflow considering only known pathogenic or likely pathogenic (LP/P) variants from humsavar as our clustering dataset. Here, VUS were more often clustered together with LP/P variants (**Supplement Figure 3**), highlighting the ability of this approach to identify functional hotspots and the possibility to connect VUS to known pathogenic variants.

### Proteorizer – an explorative tool to contextualize disease variants

As a webtool ‘Proteorizer’ makes this approach available to the scientific community. Researchers can directly input a protein and one or multiple mutations using UniProt accessions or the canonical gene name in combination with amino acid variants in IUPAC one letter code. Proteorizer currently supports only human input proteins, while alignments can consist of reviewed protein sequences of any species.

The complete set of collected information is presented in a searchable table that can be downloaded for further analysis. Additionally, Proteorizer provides a graphical representation of identified clusters and an interactive structure viewer where residues as part of relevant clusters can be highlighted. The webtool also produces an annotated alignment. This alignment displays the collected information in a predefined window (± 30 residues) around the position of interest (**Fig. 2**) but can be dynamically re-generated with different viewing ranges and different number of displayed sequences.

**Figure 2.**
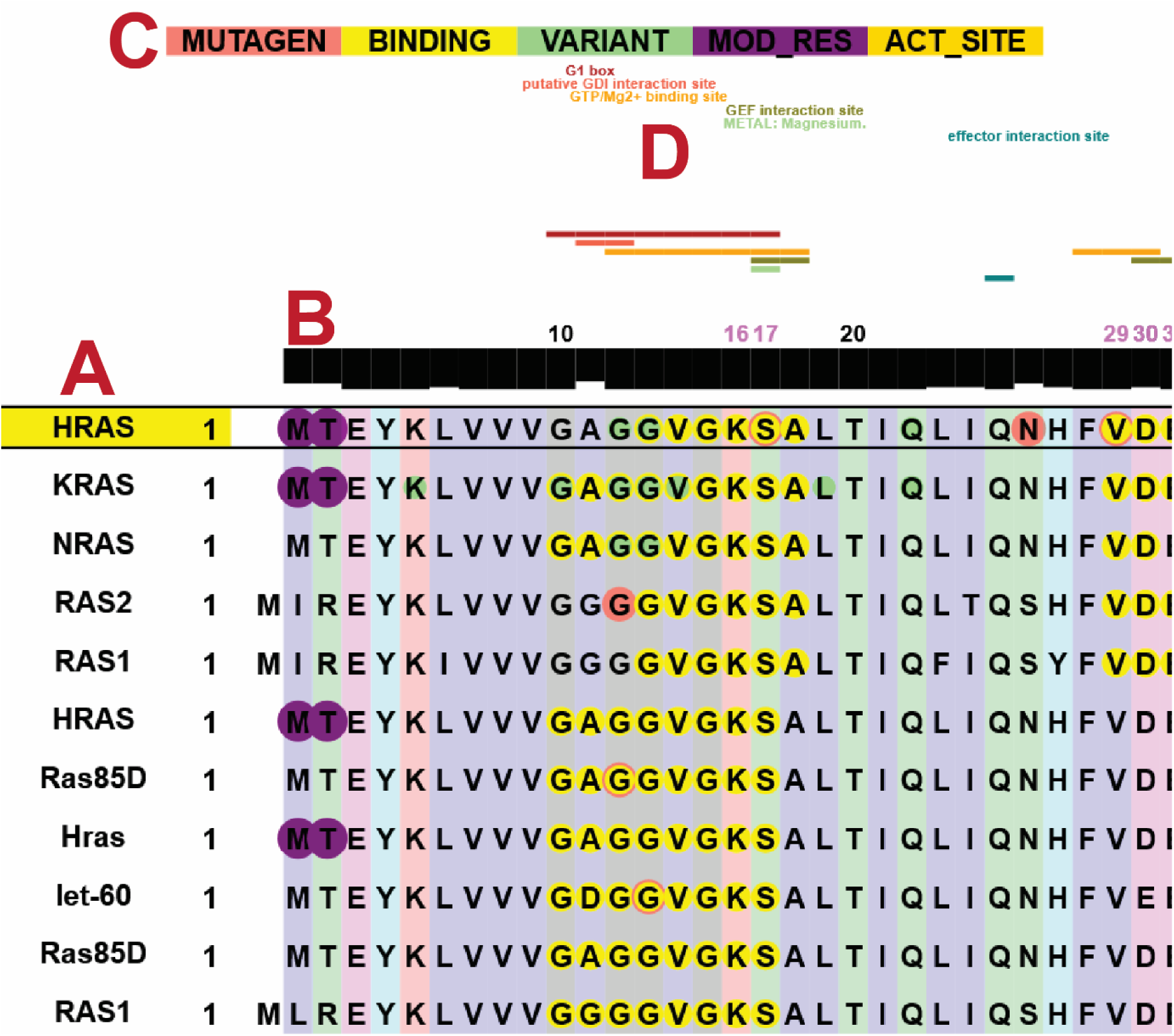
Example annotated alignment for HRAS. **A.** Gene names are given where possible (UniprotID otherwise), the gene of interest is highlighted and put at the top of the alignment. **B.** Black bars indicate the sequence identity at this position of the alignment. **C.** Residues with known functional information are highlighted with coloured circles based on the respective Uniprot categorie, ie. Salmon for mutagens, yellow for binding sites, green for observed variants, purple for PTM sites and gold for active sites. **D.** Additional protein features (such as domains or regions, taken from InterPro) are shown on top of the alignment.

### Case studies

To demonstrate different use cases of Proteorizer we applied our method using the default options on several variants previously classified as VUS.

The variant p.S244G in ALPP was seen to be exclusively heterozygous in 42 participants of the 1000 Genome Project (1kG)^30^ with additional evidence suggesting an increased chance of a functional impact^2^. We reported elsewhere that p.S244G lies close to known variants in the paralog ALPL which are associated with decreased enzyme activity^2^. Here, we additionally found that several Ca^2+^-binding sites are in close proximity, with ALPP p.Glu238 being a conserved Ca^2+^-binding site in several homologs (**Fig. 3A**). We could apply Proteorizer to add a mechanistic interpretation to a case where we previously had insufficient evidence to explain its exclusively heterozygous occurrence.

**Figure 3.**
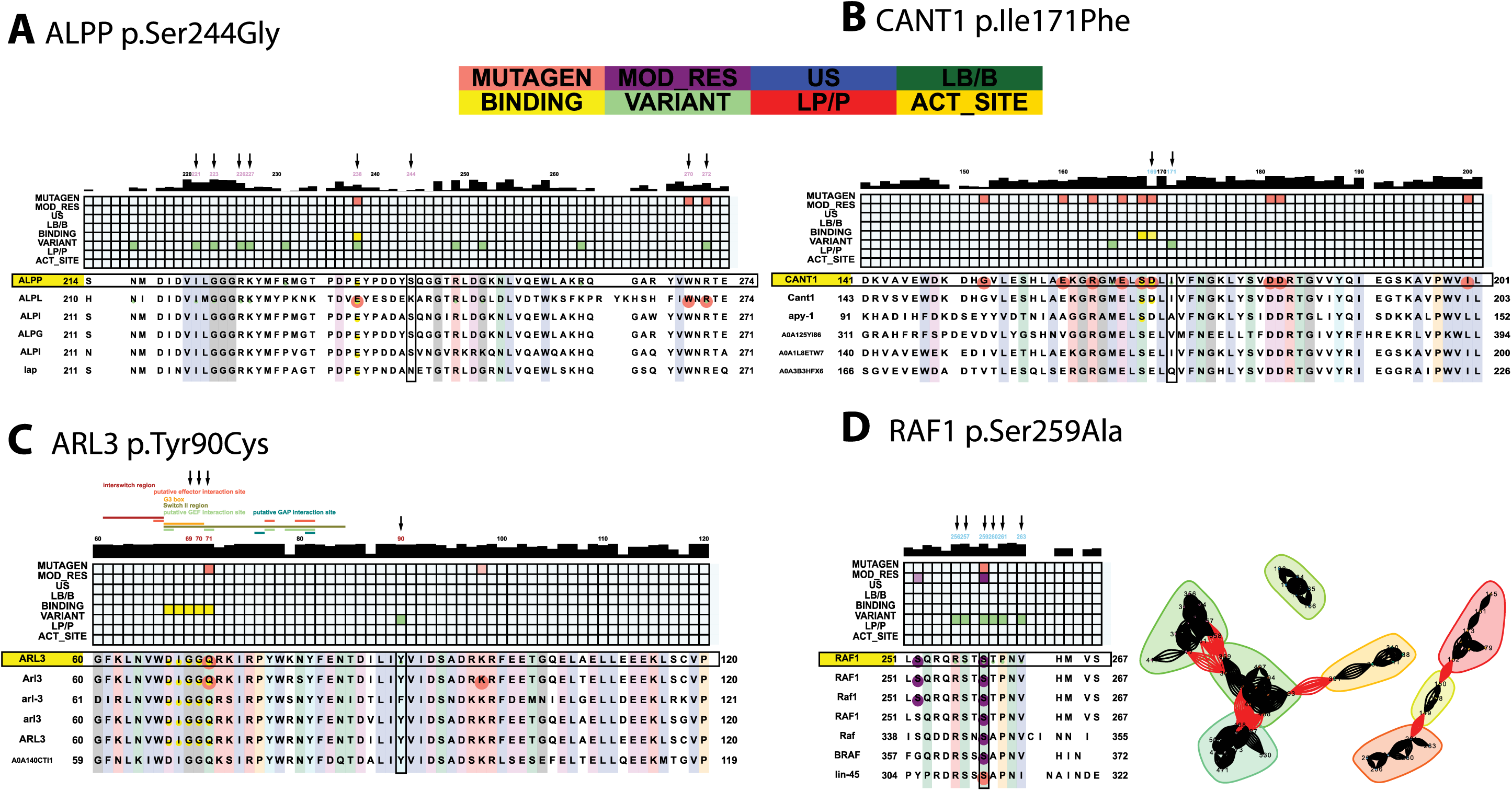
Case studies. **A.** Alignment showing functional information ± 30 positions of ALPP Ser244. Positions with functional information in the same cluster as Ser244 are highlighted with black arrows. The alignment alignment presents the collected information also as a heatmap, and positional conservation is shown above the alignment as black rectangles. **B.** As in A. shown for CANT1 p.Ile171Phe. **C.** As in A. shown for ARL3 p.Tyr90Cys. The alignment additionally also shows sequence regions with known functional annotations (Switch II etc.) above the conservational information, if available via InterPro. **D.** Left, as in A. shown for RAF1 p.Ser259Ala. Right, showing all detected functional clusters in RAF1.

The homozygous variant p.Ile171Phe in CANT1 has been found in patients affected by multiple epiphyseal dysplasia causative of abnormal skeletal development, often associated with increased morbidity^31^. CANT1 is a calcium-dependent nucleotidase likely involved in glycosylation and protein quality control^32^. The authors argue that p.Ile171Phe might result in diminished enzyme activity, similar to what has been reported for p.Val226Met^33^. Our analysis, despite not having a high quality alignment (**Fig. 3B**), indeed shows that p.Ile171Phe is located close to p.Val226Met and the Ca^2+^-binding site on position 169 (but not 168), where p.Asp169Asn was shown to reduce enzyme activity by 96%^34^. The structure-based clustering reinforces the notion that p.Ile171Phe might interfere with Ca^2+^-binding, as several other positions involved in Ca^2+^-binding (215, 284, 345 and 396) lie close (**Fig. 3B**). Our approach was able to link this VUS to a known disease variant, suggesting that p.Ile171Phe is likely causative of a similar phenotype following a comparable molecular mechanism, warranting further investigation.

The rare genetic variant p.Tyr90Cys in the small GTPase ARL3 was found in patients suffering from retinitis pigmentosa^35^, a disease associated with gradual loss of vision. Mutations in ARL3 and other Arf like proteins are associated with ciliopathies and it is hypothesized that ARL3 has functions in lipidated protein trafficking^36^. Tyr90 is highly conserved and was named a candidate for autosomal dominant retinitis pigmentosa, although the study was not able provide a mechanistic explanation^35^. Our analysis suggests that Tyr90 is fairly close to conserved residues associated with GTP binding (25, 26, 27, 28, 71) (**Fig. 3C**). Perturbation of GTP-binding is likely to decrease the activity of ARL3, and p.Gln71Leu was found to inhibit activation of mouse Arf3 via the GTPase activating protein RP2^37^. Proteorizer was able to provide a reasonable explanation using prior knowledge for a protein variant without an experimentally verified 3D structure, connecting residues that could have been easily missed by only considering a traditional 2D alignment.

Some cases might still present conflicting information. The variant p.His280Pro in IL6R has been reported as a VUS^38^. IL6R is the receptor for the cytokine IL6 and plays important roles in immune response. Although the neighbouring variant p.Ile279Asn was found to decrease STAT1 and STAT3 phosphorylation, and not p.His280Pro, our data suggests that p.His280Pro lies close to several positions with experimentally validated effects on ligand binding (232, 233, 277, 278 and 279). Interestingly, the variant p.His280Ile has no effect on ligand binding^39^. However, unlike histidine the amino acid proline is very rarely involved in protein binding events due to its very non-reactive side chain^40^. While isoleucine also has a non-reactive side chain, it has been reported that isoleucine can participate in substrate recognition events^41^. Taken together it may be possible that p.His280Pro does not ablate IL6R ligand binding and hence shows no effect on STAT1/3 phosphorylation, but the ligand binding could be obstructed enough to increase the time required for successful immune response, which can render individuals more susceptible to infectious diseases^42^. Here, the complex information presented by Proteorizer was used to offer an explanation that differs to the observations made for p.Ile279Asn on how p.His280Pro in IL6R could contribute to a disease phenotype.

Proteorizer can also be useful to generally assess whether a given variant should be investigated further. For example the variant p.Ser259Ala in RAF1 was only seen twice in COSMIC^43^, meaning the variant will typically be dismissed due to its low count. However, Proteorizer is able to assess these variants in a rapid and cost efficient manner. We can show that Ser259 is a known phosphosite not only in RAF1 but in several orthologs as well, suggesting that Ser214 in ARAF may be a phosphoserine, too (**Fig. 3D**)^44^. Additionally, Ser259 lies close to known cancer variants that increase RAF1 downstream signalling. Indeed, it was shown that Ser259 is an important inhibitory phosphosite for RAF1^20,45,46^ (**Fig. 3D**). This example highlights that our approach is a quick and inexpensive approach to prevent falsely discarding low-count variants in otherwise well-known oncogenes.

## Conclusion

Assessing the clinical relevance of VUS remains a challenge for modern precision medicine. The work presented here demonstrates how the collection of seemingly unrelated information, for example due to their perceived distance on a 2D alignment, can yield novel insights. We used this information in a holistic approach by focusing on a considerable amount of available information for the protein of interest to project this onto the 3D structure. A clustering based on intramolecular distances between residues deemed functionally interesting then produces functional hotspots. Proteorizer works as an explorative tool and we discussed our findings on several case studies. This approach combines several features into one comprehensive score and does not solely rely on sequence conservation or variant counts above a certain threshold. If a VUS is associated with a known functional hotspot it is likely to warrant further investigation.

A limitation of our approach is susceptibility to research bias, because a functional hotspot will more often be found in proteins or protein families that are already well studied, and likewise the absence of any functional hotspot could be solely due to a general lack of information (on the protein of interest or orthologs) and is therefore not necessarily an indication that a particular variant is benign. This also highlights that the predictive part of this study can only be supplemental to our hypothesis of using a wider range of prior-knowledge when investigating protein variants, and the combined score presented here should never be taken at face value. Our method is also not meant to compete with other popular predictors (Polyphen2^47^, SIFT^48^, AlphaMissense^49^) for this very reason, rather these tools should be used alongside Proteorizer.

We hope that this study can assist researchers and medical professionals alike in decision making, i.e. which experiments may be useful to conduct next, or assigning patients to promising new treatments – based on well-known residues that are part of the same functional hotspot as the respective VUS and wherever possible. The gathering, analysis and transfer of information to any user is a challenge of modern day science as well as precision medicine^50,51^, where Proteorizer aims to assist.

## Data Availability

Proteorizer is a freely accessible web tool at shiny.russelllab.org/proteorizer. The source code and additional data is available at https://github.com/tschmenger/PROTEORIZER. Other sources we used are already accessible to the public (i.e. Alphafold^5^, Interpro^52^, Humsavar^14^, Uniprot^14^).

## Code availability

Were applicable data were processed using publicly available tools described in the Methods. All scripts related to the Proteorizer web app and workflow are available via Github (see Data Availability) and a standalone script to annotate custom alignments is available via https://github.com/russelllab/Annotate_Alignments.

## Acknowledgements

We thank J.C. González-Sanchez and G. Singh for fruitful discussions and feedback.

## Author Contributions

R.B.R. and T.S. designed the study. T.S., G.D. performed the data analysis and provided the default alignments. T.S. wrote the manuscript.

## Supplementary Figures

**Supplement Figure 1.**
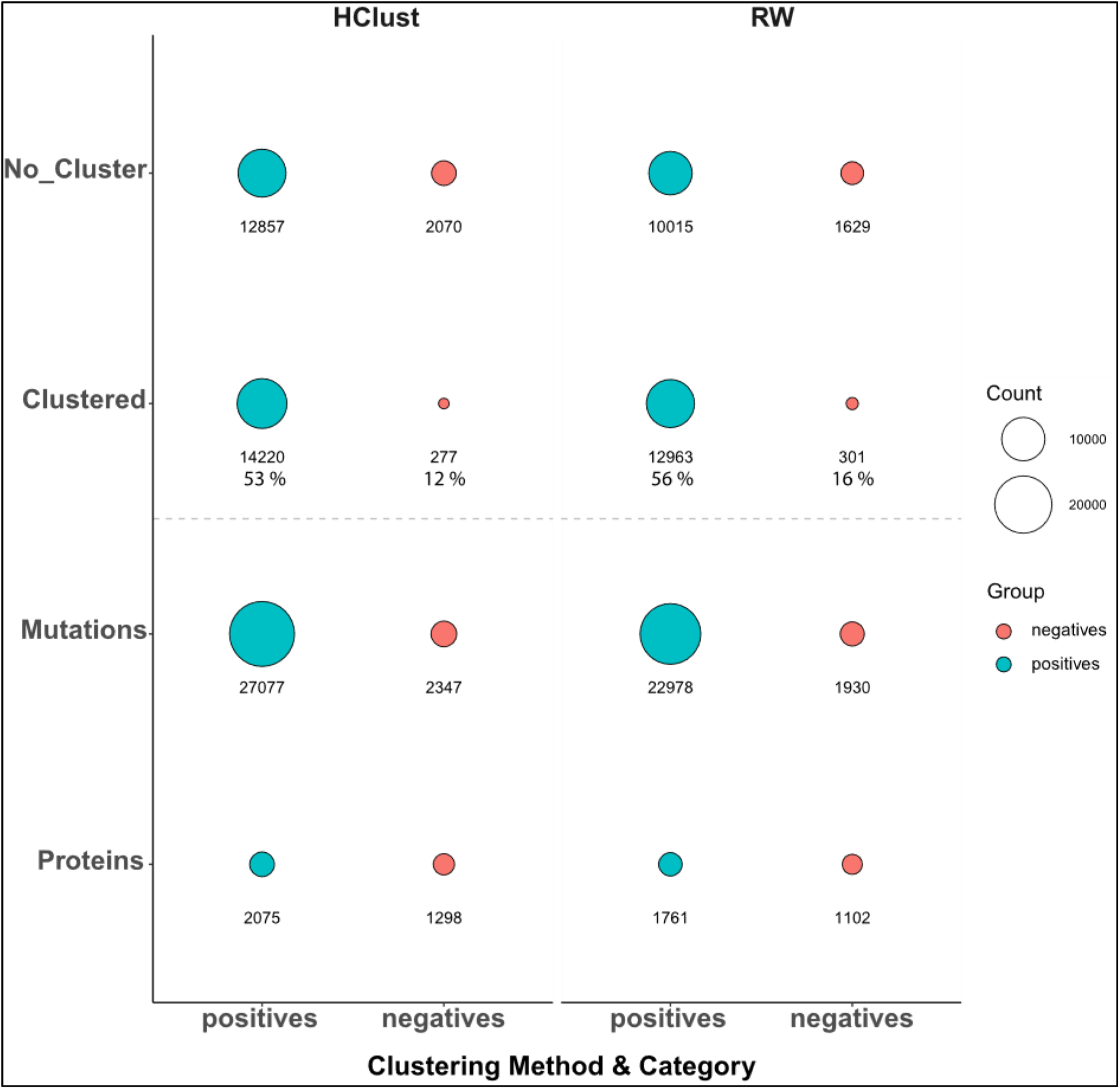
Disease variants are more likely to be part of functional clusters. Hierarchical clustering (Hclust, left) and random walk (RW, right) succeed in finding functional clusters for more than 60 % tested variants with the vast majority of clustered variants (> 50 %) being known disease variants.

**Supplement Figure 2.**
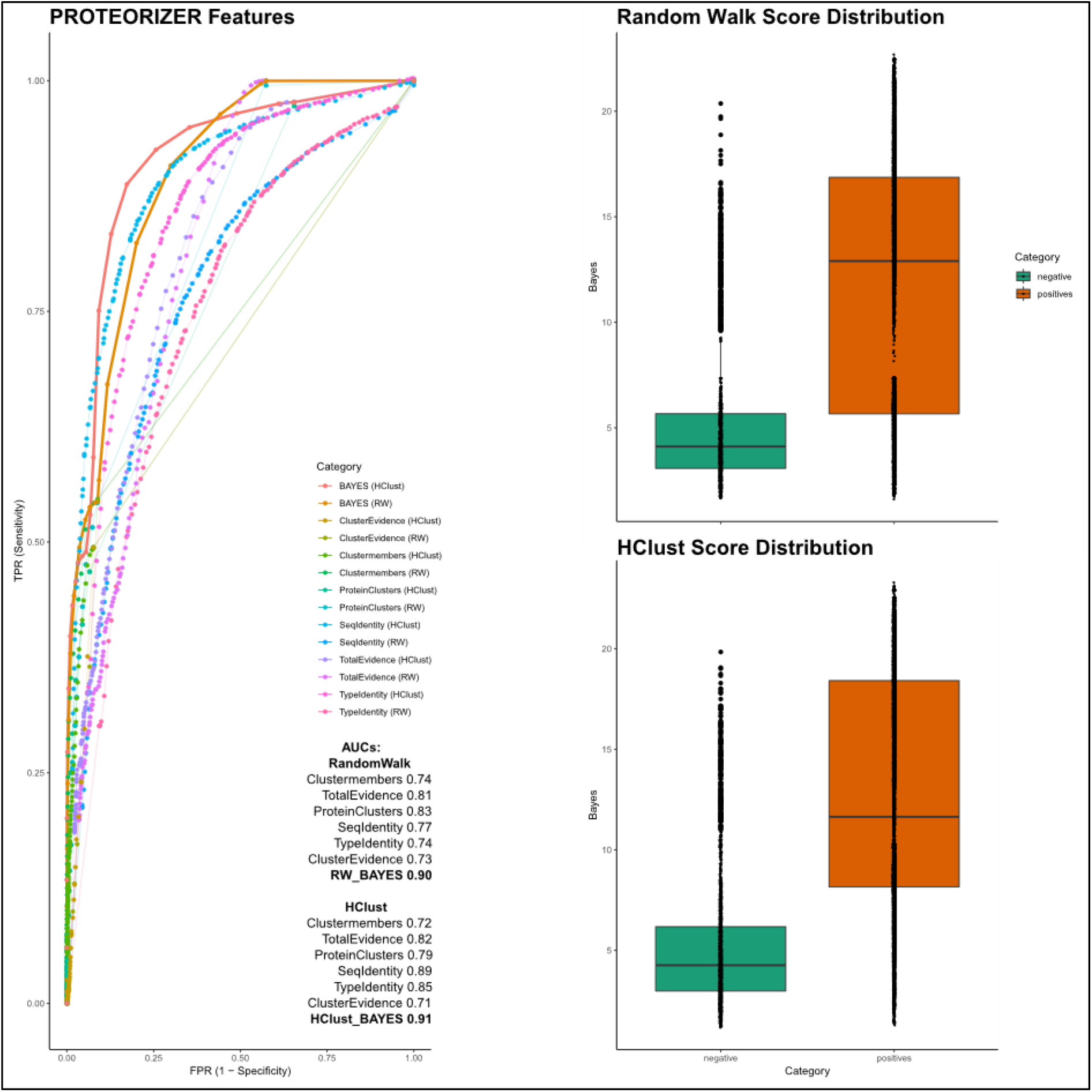
Variants as members of functional clusters as disease candidates. Bayesian combination of multiple features show good and comparable performance for both random walk and hierarchical clustering. A high Bayesian score (right top & bottom) for either hierarchical clustering or random walk suggests that a variant is likely to have a functional impact, based on features such as being part of a functional cluster, residue and information count for the respective cluster and the protein as a whole (left).

**Supplement Figure 3.**
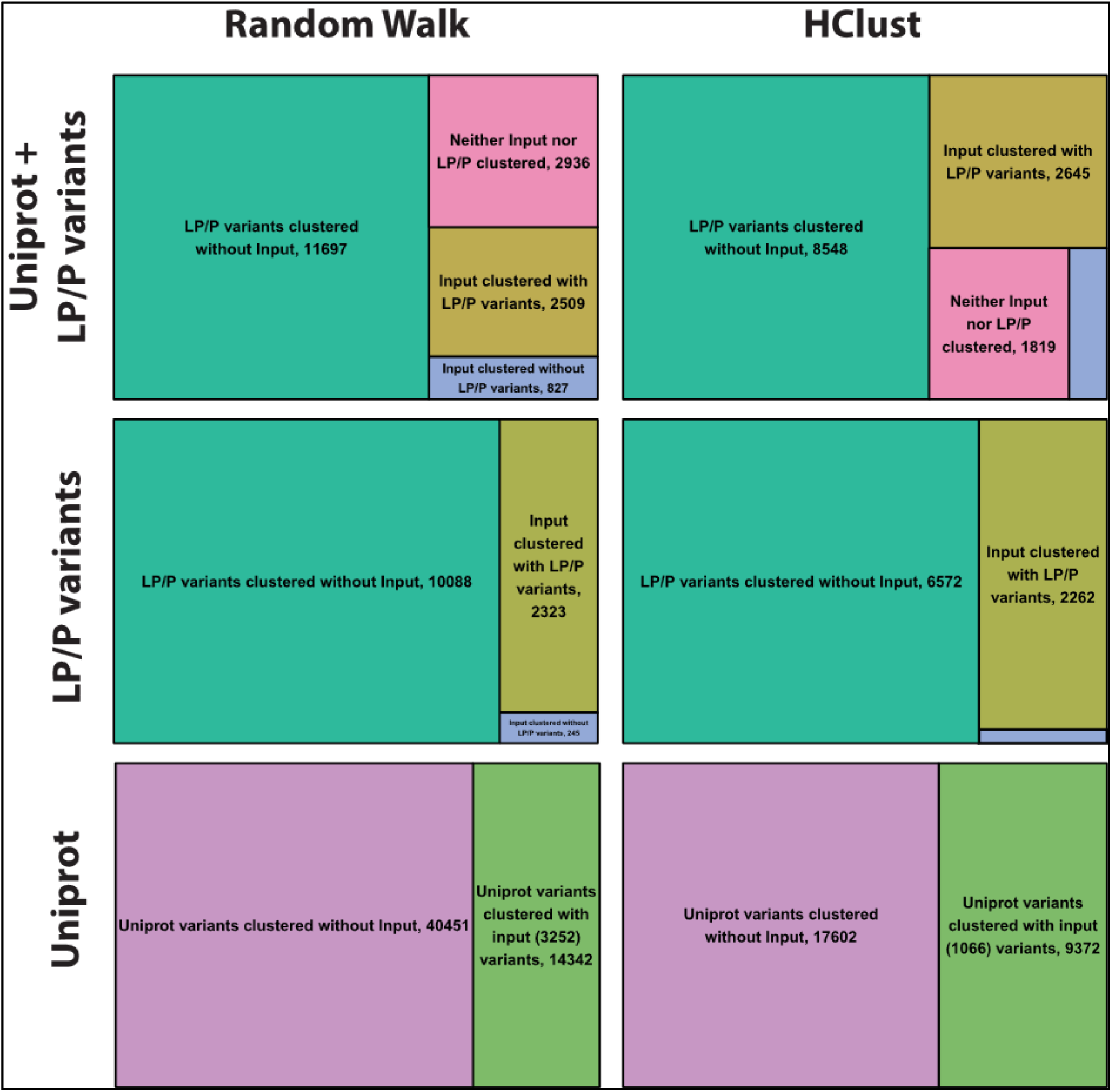
VUS cluster with LP/P more often. VUS analysed in RW or HClust mode are more often found to be clustered with known LP/P variants (top & middle, olive vs. blue).

**Supplement Figure 4.**
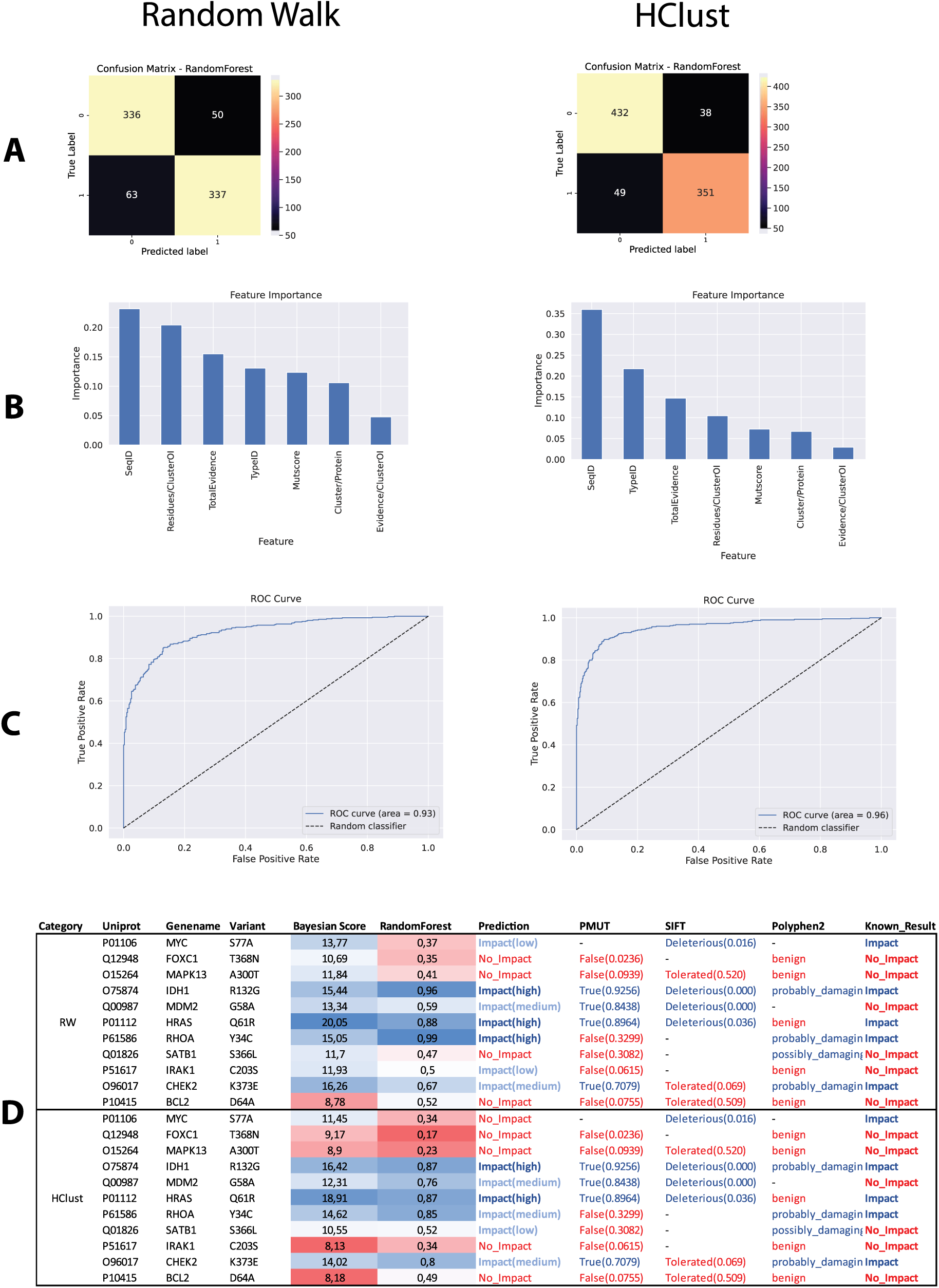
Machine-learning metrics. Left) Metrics using random walk for 3D clustering. Right) Metrics using HClust as the clustering method. **A**. Confusion matrix showing assessing the trained predictor on a subset of the training data not used during the training. **B**. Feature importance. **C**. Receiver operating curves. **D**. Additional test cases with known functional consequences that were not seen by the predictor during training.

